# Quantitative Protein Topography Measurements by High Resolution Hydroxyl Radical Protein Footprinting Enable Accurate Molecular Model Selection

**DOI:** 10.1101/136929

**Authors:** Boer Xie, Amika Sood, Robert J. Woods, Joshua S. Sharp

**Author notes:** Correspondence to: Joshua S. Sharp; Department of BioMolecular Sciences P.O. Box 1848, University, MS 38677-1848. These authors contributed equally to this work (Boer Xie, Amika Sood).

## Abstract

We report an integrated workflow that allows mass spectrometry-based high-resolution hydroxyl radical protein footprinting (HR-HRPF) measurements to accurately measure the absolute average solvent accessible surface area (<SASA>) of amino acid side chains. This approach is based on application of multi-point HR-HRPF, electron-transfer dissociation (ETD) tandem MS (MS/MS) acquisition, measurement of effective radical doses by radical dosimetry, and proper normalization of the inherent reactivity of the amino acids. The accuracy of the resulting <SASA> measurements was tested by using well-characterized protein models. Moreover, we demonstrated the ability to use <SASA> measurements from HR-HRPF to differentiate molecular models of high accuracy (< 3Å backbone RMSD) from models of lower accuracy (> 4Å backbone RMSD). The ability of <SASA> data from HR-HRPF to differentiate molecular model quality was found to be comparable to that of <SASA> data obtained from X-ray crystal structures, indicating the accuracy and utility of HR-HRPF for evaluating the accuracy of computational models.

## INTRODUCTION

The generation of accurate high resolution models has revolutionized protein structure-function studies, but the determination of protein structure is often the most difficult and laborious step. Many important protein systems have proven difficult to interrogate by traditional high resolution structural biology methods due to difficulties in crystallization, requirements for sample purity and amount, and other experimental difficulties with high resolution structural biology methods. Mass spectrometry (MS)-based structural methods have proven quite valuable in such cases,^1^-^5^ however, they usually do not provide structural data of sufficient quantitative accuracy and structural resolution to be employed as constraints or quantitative metrics for high resolution structural modeling. A need exists for technologies that can address the challenging protein targets, while generating data of sufficient resolution and accuracy to base molecular modeling efforts around.

One method that has been used with considerable success to study changes in protein topography is hydroxyl radical protein footprinting (HRPF). HRPF entails oxidizing amino acid side chains in a protein of interest with diffusing hydroxyl radicals generated *in situ* by a variety of means, including UV laser-based flash photolysis of hydrogen peroxide.^*^6^*^ These hydroxyl radicals are broadly reactive and mimic water in size and polarity, labeling amino acids on the surface of the folded protein structure.^2, 6-9^ The apparent rate of hydroxyl radical-mediated oxidation of a given stretch of amino acids depends on the inherent reactivity of that portion of the protein to hydroxyl radicals, as well as the accessibility of the segment to the radical as it diffused through the solvent.^10^ Until recently, oxidation was quantified at the peptide level by liquid chromatography coupled to mass spectrometry (LC-MS), allowing for measurements of changes in protein topography to be made at a moderate structural resolution, limited by the size of the proteolytic peptide being measured.^11, 12^

More recently, a variety of new methods in HRPF allow for high resolution maps of changes in protein topography to be made with many amino acids probed in a single experiment. The development of the widely used Fast Photochemical Oxidation of Proteins (FPOP) technology for HRPF experiments allows for more thorough oxidation of target proteins while maintaining their native conformation,^6, 13-15^ resulting in multiple modification sites within a single peptide sequence and a higher density of structural information.^6, 13, 15^ Methods allowing accurate quantification of multiple sites of oxidation spread out along a single peptide allow for the higher density of oxidation available in FPOP experiments to be properly measured,^14, 16-18^ and new methods for monitoring the effects of radical generation and scavenging on the effective concentration of available radical now allow for normalization of different experimental conditions.^19, 20^ While this combination of new technologies for HRPF now allows for higher resolution monitoring of changes in protein topography, there is currently no method for converting HRPF data into a quantitative measure of absolute protein topography; data are reported as relative changes between two protein conformations (e.g. ligand-bound versus ligand-free). If accurate absolute <SASA> data were available, molecular models could be evaluated quantitatively for agreement with experimental results, potentially offering a new MS-based method for generating accurate, experimentally validated molecular models of protein structure.

Since the earliest days of MS-based HRPF, it has been observed that the apparent rate of hydroxyl radical oxidation of a peptide or protein correlated with the solvent accessibility of that protein.^21^ Previous reports by Charvatova *et al.* have shown a quantitative correlation between HRPF data and <SASA> of constituent amino acids,^22^ and a later report from Huang et al.^23^ explored the possibility of using peptide-resolution HRPF to derive HRPF-based protection factors. However, these studies largely depended upon peptide-level resolution HRPF data. Moreover, they assumed that all amino acids of a particular structure had the same inherent reactivity as previously measured for free amino acids,^24^ which was brought into question in previous studies of model peptides.^25^ The observed correlation was insufficient for accurate determinations of <SASA> based on HRPF data. The correlation between LC-based high resolution HRPF data using FPOP of barstar also showed insufficient correlation between HRPF data and <SASA>.^14^

Here, we introduce a method for generating accurate side chain absolute <SASA> values for a wide variety of amino acids using FPOP HR-HRPF data by integrating a variety of technological improvements for FPOP previously described by our group, as well as by developing improved methods to normalize for inherent amino acid reactivity. Using the technologies described here, we present data using the model protein system lysozyme demonstrating that we can clearly differentiate between high-quality molecular models (< 3Å backbone RMSD) and models of lower accuracy (> 4Å backbone RMSD) based on the agreement of the model’s <SASA> values with those measured experimentally using HR-HRPF. With these advances, the generation of accurate molecular models of protein structure validated by MS HR-HRPF data becomes truly possible.

## RESULTS

We sought to design an integrated workflow to convert HR-HRPF data into quantitative measurements of protein topography. In order to meet this goal, several obstacles needed to be addressed: 1.) HR-HRPF apparent oxidation rates must be accurately measured at the amino acid level; 2.) the measured oxidation must be normalized by the concentration of radical generated and the scavenging properties of the solution; 3.) the inherent reactivity of the amino acid *in its sequence context* must be accurately measured or estimated; 4.) the quantitative relationship between normalized amino acid reactivity and <SASA> must be established.

### Multi-point HR-HRPF analysis

Accurately measuring the apparent rate of HR-HRPF oxidation by FPOP is key to any experimental attempt to link HR-HRPF measurements to absolute biophysical properties of the protein. Other HRPF methods, most notably synchrotron radiolysis methods, have long used multi-point HRPF analysis to improve accuracy and ensure the HRPF labeling process does not unfold the protein.^1^ However, due to the observed ability of FPOP to label proteins faster than label-induced conformational changes can occur,^15^ multi-point FPOP is not typically employed. We used multi-point FPOP HR-HRPF measurements to improve the measurement accuracy of apparent rate of reaction measurements, altering the concentration of hydrogen peroxide exposed to the UV laser to alter the concentration of radical generated. Due to the broad reactivity of the hydroxyl radical, controlling the amount of radical *generated* is not sufficient; the amount of radical consumed by radical scavengers in solution (e.g. buffers, carrier proteins, etc.) must also be determined in order to understand the *effective radical dose* experienced by the protein (which is dependent upon the radical concentration and the radical half-life). ETD-based LC-MS/MS analyses are performed to accurately measure oxidation of individual amino acid side chains.^17, 18^ We employ an adenine radical dosimeter^20^ to measure the effective radical dose under different experimental conditions, and plot HR-HRPF oxidation based on the adenine dosimeter response in Figure 1. The measured reactivity from the regression slope reflects the radical response rate of an oxidized amino acid.

**Figure 1.**
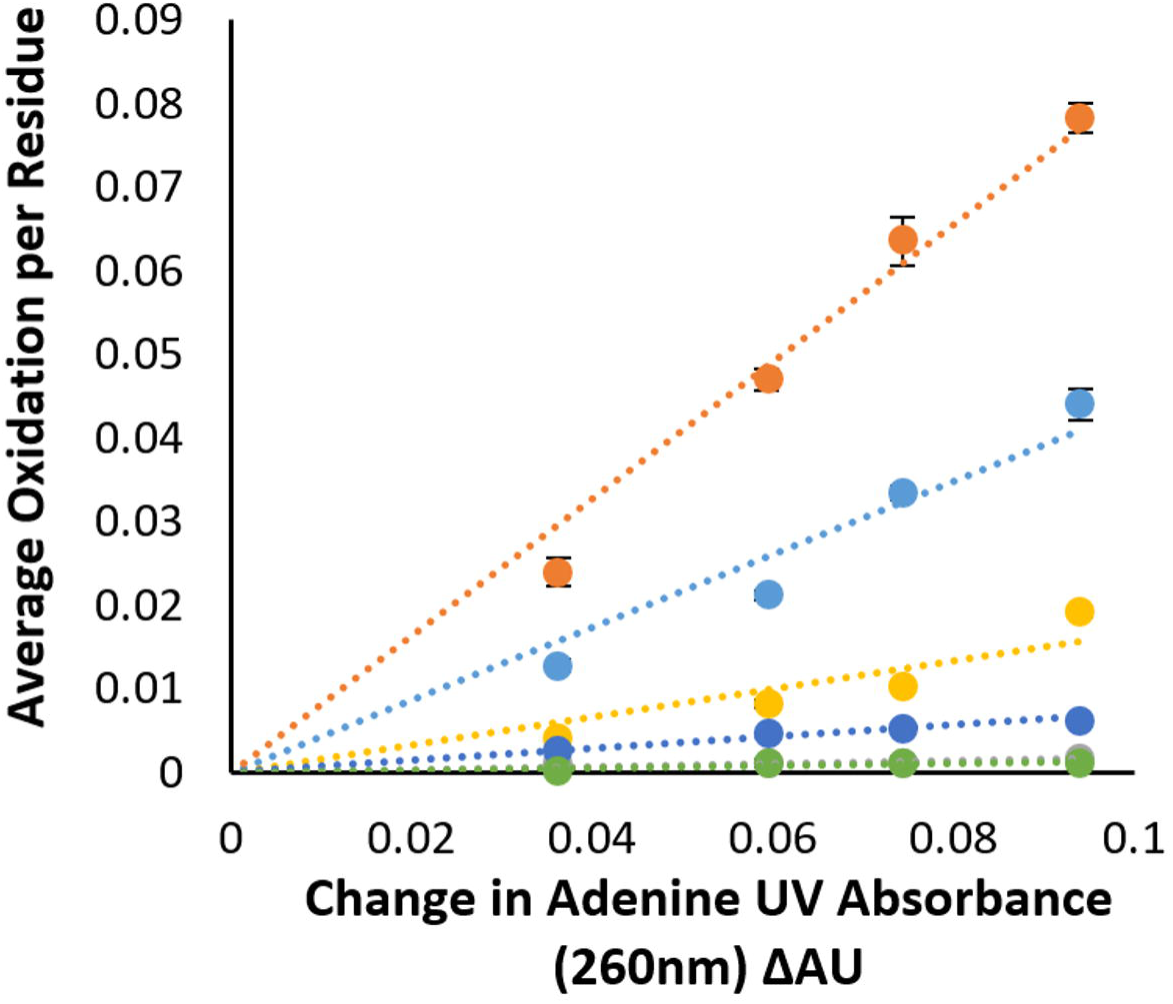
Representative HR-HRPF regressions for five residues from lysozyme after multi-point HR-HRPF performed in triplicate (mean ± SD plotted; error bars obscured by data markers). The levels of oxidation vary linearly with the available hydroxyl radical dose, measured by adenine dosimetry; reactivity is measured as the slope of the regression. Orange - Trp123, light blue – Trp111, yellow – Phe34, dark blue – Asn37, and green – Asn39.

The radical response rate, which reflect the reactivity of a given residue in a given protein sequence and a given conformation to hydroxyl radicals in the HR-HRPF experiment, is a function of the intrinsic reactivity of the residue to hydroxyl radical, as well as its steric availability to attack by the hydroxyl radical, estimated by its <SASA>.^21-23^ In practice, a free amino acid has a specific intrinsic reactivity to hydroxyl radicals that vary from ∼10^7^ to 10^10^ s^-1^.^24, 26^ Because of this broad range of reactivity, the intrinsic reactivity becomes an important factor for characterization of solvent accessibility at different sites using the radical response rate, and has historically been normalized based on the reactivity of the free amino acid which generate a normalized protection factor (NPF) that reflects radical response rate of a specific residue *R*_*i*_.^23^ However, little work has been reported to indicate if the intrinsic reactivity of a free amino acid is an accurate representation of the intrinsic reactivity of that amino acid within a polypeptide sequence context. Indeed, the one study examining the question suggests that, at least for some amino acids in some contexts, sequence context can change the apparent rate of oxidation significantly.^25^

To explore the extent by which radical response rates normalized by free amino acid inherent reactivities capture protein structural information, we examined the correlation between NPF from multi-point HR-FRPF data and fractional <SASA>, which is obtained by comparing the <SASA> of the amino acid in the folded structure against the fully-exposed <SASA> as calculated from a Gly-X-Gly peptide (Figure 2). Different free amino acids have widely different <SASA> values, and bulkier amino acids will sterically encounter radicals more often than smaller amino acids (e.g. a large factor in the higher reactivity of Leu compared to Val is simply the fact that Leu is larger and has more C-H bonds for the radical to attack). We calculated the fractional exposure of each amino acid in the native structure by dividing the <SASA> of the amino acid within the MD trajectory of the folded structure by the <SASA> of that amino acid in a Gly-X-Gly peptide, as previously reported.^27^ While we believe this normalization model better reflects the biophysical realities underlying reactivity, in no case here did it make a substantial difference in the data presented (data not shown). The NPFs of two model proteins, lysozyme and myoglobin, were measured by multi-point HR-HRPF analysis. As previously observed, the apparent reactivity of the same type of amino acid side chain can vary widely based on the <SASA>. Higher <SASA> tends to give more oxidation for amino acids of similar inherent reactivity; e.g. Trp123 (<SASA>=71 Å^2^) shows a much higher reactivity than Trp111 (<SASA>=25 Å^2^) (Fig. 1).

**Figure 2.**
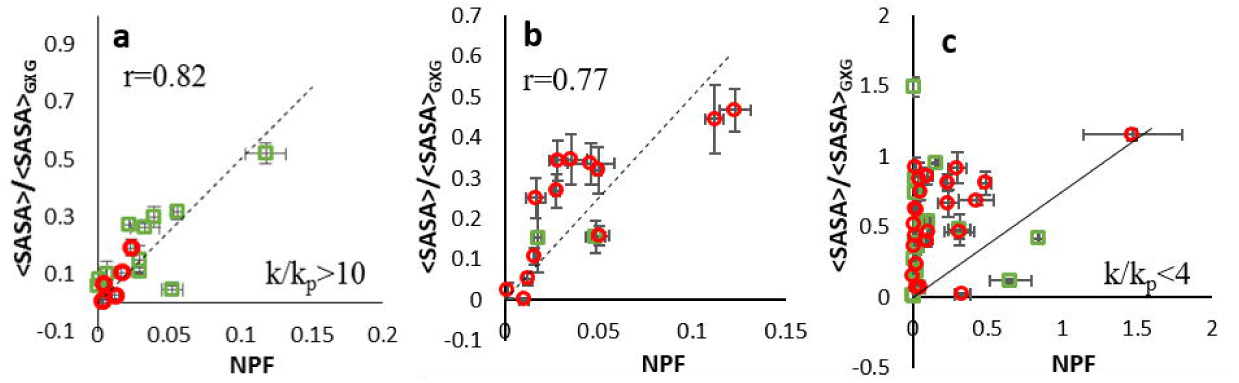
Correlation plots of NPF (the slope of the linear regression divided by the inherent reactivity of the free amino acid) vs. fractional <SASA> (<SASA> of the amino acid in the folded structure divided by the <SASA> for that same amino acid in a Gly-X-Gly tripeptide) for amino acids identified to be oxidized from lysozyme (green square) and myoglobin (red circle) sample for: (**a**) amino acids with high inherent reactivity (>10x more reactive than Pro); (**b**) amino acids with moderate inherent reactivity (4-10x more reactive than Pro); (**c**) amino acids with poor inherent reactivity (<4x more reactive than Pro). All data shown from triplicate experiments using mean ± SD plotted for all data (x-axis error bars), with fractional <SASA> from MD simulation of the protein structure ± SD from the MD trajectory (y-axis error bars).

A total of 72 amino acids were identified to be oxidized from the FPOP labeling: 30 amino acids from lysozyme, and 42 from myoglobin (**Supplementary Table 1**). The sulfur-containing residues were excluded from the correlation studies due to their extremely high reactivity to reactive oxygen species of all kinds and their susceptibility to background oxidation before, during, and after oxidation, resulting in artificially high levels of oxidation after normalization when compared to other amino acids of similar inherent reactivity.^2, 16, 28-30^ We divided the remaining amino acids based on the reactivity of the free amino acid compared to Pro: highly reactive residues (k/k_p_>10, Fig. 2a), moderately reactive residues (4<k/k_p_<10, Fig. 2b), and poorly reactive residues (k/k_p_<4, Fig. 2c).^24^ In the case of highly reactive residues with k/k_p_>10, a strong correlation can be observed between measured NPFs and fractional <SASA> with Pearson correlation coefficient (r) of 0.82 suggesting that the model relying upon free amino acid reactivities works reasonably well. The moderately reactive residues show a slightly worse correlation (r=0.77). The poorly reactive residues show a statistically insignificant correlation (r=0.24), even though the confidence and precision of the HR-HRPF measurements remained high for almost all data points. These data clearly indicate that normalization of amino acid HR-HRPF by free amino acid reactivity loses accuracy as the inherent reactivity of the amino acid decreases. However, previous studies looking at changes of <SASA> of amino acids with poor free amino acid reactivity upon ligand binding were able to successfully identify the ligand binding sites,^18^ indicating that the reactivity of these poorly reactive residues is still impacted by <SASA>. It is possible that the sequence context of a folded native protein has a significant effect on the inherent reactivity of each amino acid, and that the relative impact of the sequence context is greater for amino acids with lower inherent reactivity. This is consistent with a previous study examining the effect of sequence context on hydroxyl radical oxidation in short model peptides, which focused on the oxidation of Leu (a poorly-reactive amino acid) and found a sizable effect of sequence context on oxdiation.^25^ If sequence context does play a significant role in the inherent reactivity of amino acids, a method for correcting for this sequence context must be developed for poorly reactive amino acids to be useful for absolute topographical analysis.

In order to test if sequence context explains the majority of the deviations we observe in correlating <SASA> to multi-point HR-HRPF measurements, we tested our ability to predict <SASA> from multi-point HR-HRPF analysis without relying upon free amino acid reactivities for normalization. We instead compared the measured reactivity of each amino acid in a native fold to the measured reactivity of that same amino acid in a thermally denatured protein, minimizing the contact of the side chains from neighboring residues while maintaining the sequence context. The denatured protein data showed appearance of oxidation on residues with high intrinsic reactivity to hydroxyl radicals but low <SASA> in the native form, as well as disappearance of oxidation on poorly reactive residues that were oxidized in the native form due to increased competition from other residues. A total of 24 amino acids from myoglobin and 16 from lysozyme were found to be oxidized in both the native and denatured conformations (**Supplementary Table 2**). Plotting the ratio of the reactivity (native: denatured) versus the ratio of <SASA> (native: denatured) greatly improves the correlation between measured radical response rates from multi-point HR-HRPF data and fractional <SASA> (Figure 3): r=0.85 for all amino acids measured, with no noticeable change in correlation between highly reactive (Fig. 3, red) and poorly reactive (Fig. 3, green) amino acids. Correlating the fractional <SASA> of the same set of amino acids using multi-point HR-HRPF data from the native fold and normalizing by free amino acid reactivity results in a much worse correlation with a wide deviation of poorly reactive residues (**Supplementary Fig. S1**). These results clearly indicate that the sequence context of a folded protein can significantly effect on the inherent reactivity of each amino acid, and this effect accounts for the bulk of the error found in correlations attempting to normalize based on free amino acid reactivities, especially for less reactive residues. In order to build a robust model for accurate quantitative topographical measurements by HR-HRPF, sequence context effect should be taken into consideration into HR-HRPF analysis.

**Figure 3.**
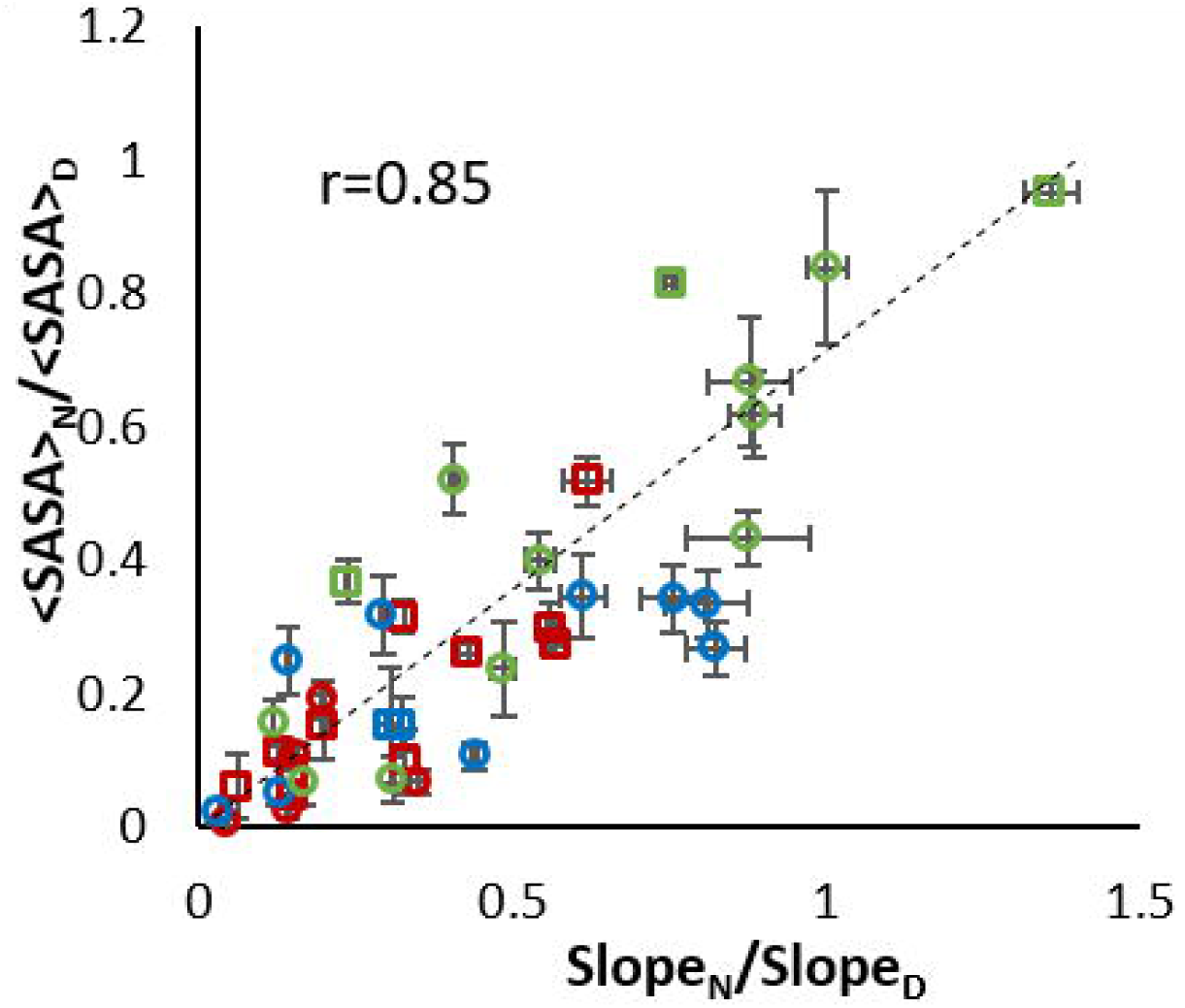
Correlation between multi-point HR-HRPF data by comparing reactivities in denatured form (Slope_D_) versus native form (Slope_N_) and fractional <SASA> between native (<SASA>_N_) and denatured (<SASA>_D_) based on all non-sulfur containing residues from myoglobin (circles) and lysozyme (squares). Each data point represents an oxidized amino acid identified in both denatured and native sample. Red – highly reactive residues; Blue – moderately reactive residues; and Green – poorly reactive residues. Data shown from triplicate experiments using mean ± SD plotted for all data (x-axis error bars), with fractional <SASA> from MD simulation of the protein structure ± SD from the MD trajectory (y-axis error bars).

A similar approach was used to test if the secondary structure context of an amino acid affects its measured HR-HRPF reactivity. Shown in **Supplementary Fig. S2** are the data from Figure 3 grouped by secondary structure element, as assigned in the X-ray crystal structures (PDB ID 1YMB and 2LYZ). The incorporation of an amino acid in a secondary structural element has no clear effect on either the magnitude or the direction of deviation of the HR-HRPF relationship with <SASA>. This indicates that secondary structural context has little to no effect of HR-HRPF reactivity, outside of its direct effect on <SASA>.

### Accuracy of FPOP multi-point reactivity analysis on predicting <SASA>

To assess the use of multi-point reactivity analysis normalized by free amino acids for estimating <SASA>, we developed empirical <SASA> prediction models based on the linear regression obtained from the multi-point HR-HRPF data for myoglobin, and a second model made from the combination of HR-HRPF data from both proteins. The linear regression models were derived based on multi-point HR-HRPF data from myoglobin sample only (single protein-based linear regression model) or from both myoglobin and lysozyme samples (dual protein-based linear regression model) using all the non-sulfur containing residues with k/k_p_>4. All residues with k/k_p_<4 were excluded from model due to their poorly correlation with <SASA> based on free amino acid reactivity normalization. The proposed empirical models were then tested for their ability to successfully predict <SASA> (as calculated from an MD simulation based on the X-ray crystal structure) using multi-point HR-HRPF data from lysozyme. Similar correlations and predictive models were observed for both single protein-based model (r=0.82, Fig. 4a) and dual protein-based model (r=0.80, **Supplementary Fig. S3a**). The robustness of these two models were tested using a leave-one-out jackknife test. Statistically significant and robust correlations were observed between measured and MD simulation <SASA> (**Supplementary Fig. S4**). Both empirical models were able to provide accurate estimated lysozyme <SASA> comparable to the actual lysozyme <SASA> from MD simulation, and providing <SASA> root-mean-square deviation (RMSD) of 19.4 Å^2^ for single protein-based linear regression model (R^2^ = 0.62) and 19.3 Å^2^ (R^2^ = 0.63) for dual protein-based model (Fig. 4b and **Supplementary Fig. S3b**). The R^2^ value and RMSD results suggest that both proposed models were able to predict <SASA> directly from multi-point HR-HRPF data with similar accuracy. Since the addition of the redundant lysozyme data to the model does not markedly improve the <SASA> measurement of lysozyme itself over the myoglobin-only model, the myoglobin-only model is a suitable empirical model for evaluating the unrelated lysozyme HRPF data, and therefore probably a suitable model for evaluating globular protein <SASA> data in general.

**Figure 4.**
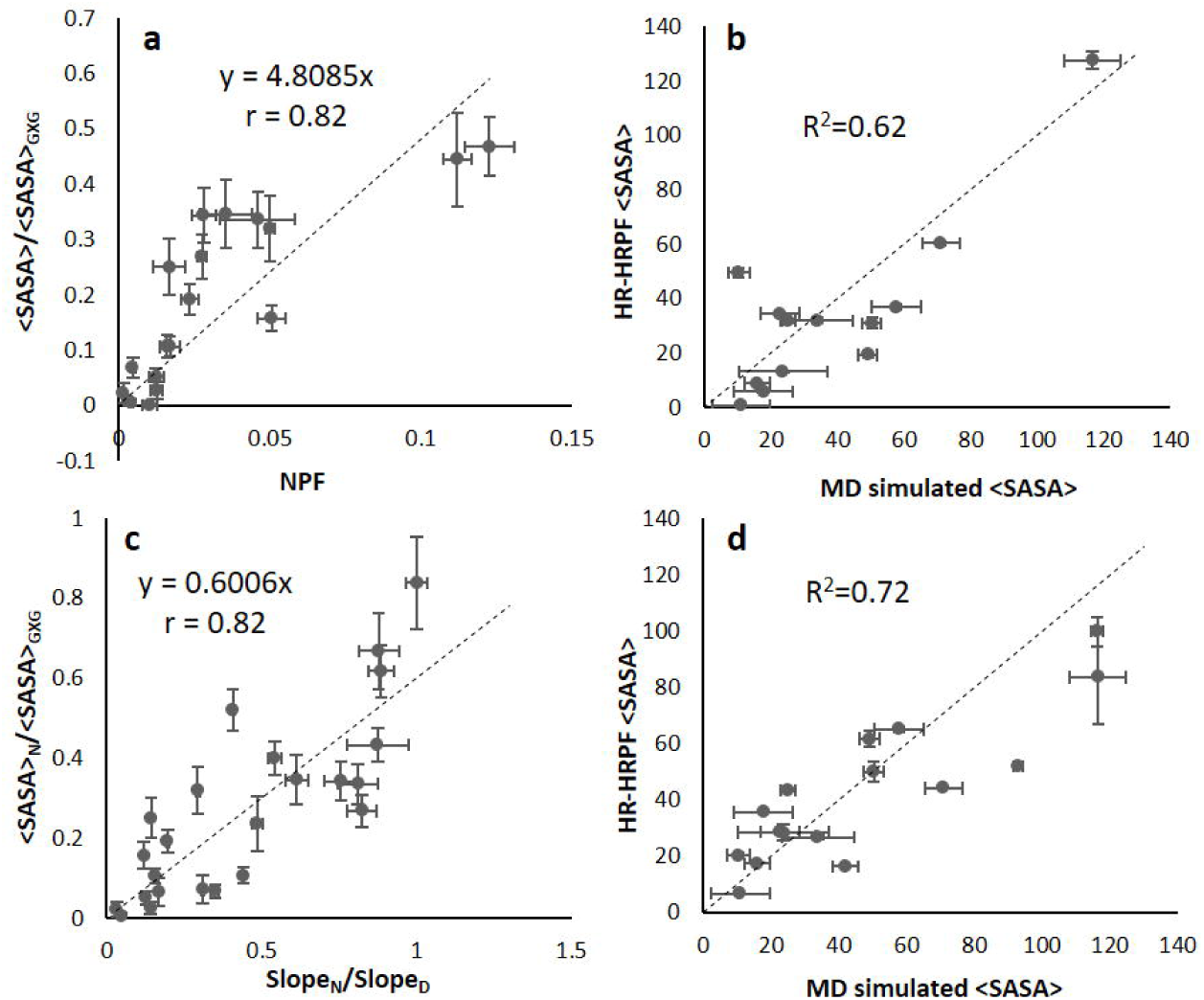
Evaluation of <SASA> prediction regression models for HR-HRPF data. (**a**) Predictive model based on multi-point HR-HRPF data from all non-sulfur containing residues with k/ k_p_ > 4 from myoglobin based on NPFs calculated from multi-point HR-HRPF data. (**b**) Comparison between predicted lysozyme <SASA> by HR-HRPF and actual lysozyme <SASA> from MD simulation based on NPFs calculated from multi-point HR-HRPF data. Correlation coefficient calculated against a regression where y=x. (**c**) Predictive model based on multi-point HR-HRPF data from all non-sulfur containing residues from myoglobin based on the ratio of reactivities (native: denatured). (**d**) Comparison between predicted lysozyme <SASA> by HRPF and actual lysozyme <SASA> from MD simulation based on the ratio of reactivities (native: denatured). Mean ± SD plotted for all data from triplicate experiments.

The proposed empirical models provide the possibility for reasonably accurate prediction of <SASA> for residues with k/k_p_>4. However, a substantial loss in both accuracy and robustness was observed when including less reactive residues (**Supplementary Fig. S5**). The MD simulation <SASA> values for the aromatic and aliphatic amino acids, which includes most of the amino acids with k/k_p_>4, were found to correlate most strongly with the backbone RMSD accuracy, indicating that these residues are the most useful for assessing the accuracy of theoretical models (**Supplementary Fig. S6**). Thus, the accuracy of the empirical model obtained from FPOP multi-point reactivity analysis normalized by free amino acids reactivity when predicting <SASA> depends heavily on the residues included in the model, and accuracy and robustness can be achieved when less reactive residues are excluded. However, sufficient divergence from the model exists to suggest that, even for the highly reactive residues, there are other factors unaccounted for in our model affecting reactivity.

Since the sequence context effects play a more prominent role for less reactive residues, the most reliable estimation of <SASA> from multi-point HR-HRPF data can be achieved by comparing HRPF reactivities in a fully denatured form versus the native folded form. By using this method, we investigated the correlations of the ratio of reactivity (native: denatured) with the ratio of <SASA> (native: denatured) and proposed two regression models using same standard proteins as for the native fold studies. This approach assumes the <SASA> of a residue in the denatured protein can be accurately modeled based on the <SASA> of that residue in a Gly-X-Gly tripeptide. A good correlation can be observed for both single protein-based linear regression model (based on multi-point HR-HRPF data from myoglobin only) and dual protein-based linear regression model (based on multi-point HR-HRPF data from both myoglobin and lysozyme) with Pearson correlation coefficient of 0.82 and 0.85, respectively (Fig. 4c and **Supplementary Fig. S7a**). The leave-one-out jackknife test was also performed on the two proposed regression models to test their robustness. Again, statistically significant correlations were observed between predicted fractional <SASA> and actual fractional <SASA> derived from MD simulation. A prefect overlapping of jackknife model and overall model indicates the robustness of the two proposed empirical models (**Supplementary Fig. S8**).

The ability to accurately derive <SASA> using the two proposed empirical models derived from multi-point HR-HRPF by comparative measurement of the conformation with a fully denatured protein were tested against the <SASA> calculated from MD simulation. Both single protein-based linear regression model (R^2^=0.72, RMSD=19.24 Å^2^, Fig. 4d) and dual protein-based linear regression model (R^2^=0.73, RMSD=17.66 Å^2^, **Supplementary Fig. S7b**) were able to provide sufficient accuracy for estimating lysozyme <SASA> as compared to the MD simulation derived lysozyme <SASA>. An improved R^2^ was observed for both models compared to the previous approach based upon the inherent reactivity measurements of free amino acids. This indicates the model based on the multi-point HR-HRPF data using ratio of reactivity (native: denatured) improves the fit of the model, which should be able to provide a better prediction of <SASA>. The number of data points used to build the model has an insignificant effect on the overall <SASA> estimation accuracy regardless of the multi-point HR-HRPF analysis method used to derive the regression model. However, the RMSD indicates a comparable accuracy for <SASA> prediction by either the model based upon the inherent reactivity measurements of free amino acids or the model based on multi-point HR-HRPF data by comparing reactivities in denatured form versus native form, the previous attempt depends heavily on the residues included in the model whereas the later approach does not influence by the residue types used to derive the model.

### HR-HRPF derived SASA RMSD for assessing the quality of protein structural models

To evaluate the performance of HRPF derived <SASA> as quantitative metrics for structural modeling, the SASA RMSD scores were calculated using the data derived from the single protein-based linear regression models from each approach proposed in this study. An unfolding simulation of the globular protein lysozyme was performed and the SASA RMSD values were calculated using SASA_fold_ values derived from the crystal structure, and from HR-HRPF data. As the protein unfolds, the SASA RMSD increases (Figure 5 a-b). The similarity of the SASA RMSD values computed using the crystallographic SASA and HR-HRPF <SASA> references reflects the excellent agreement between the <SASA> from the HR-HRPF experiments and those from the crystallographic data. A better overlapping of HR-HRPF derived <SASA> with crystallographic SASA, especially at lower RMSD range, was observed by the approach using the ratio of reactivity (native: denatured). The HR-HRPF derived <SASA> from both approaches appear to be suitable to use as quantitative metrics for model curation, with the ratio of reactivity (native: denatured) data proving slightly more accurate and better able to discriminate between bad and good structures.

**Figure 5.**
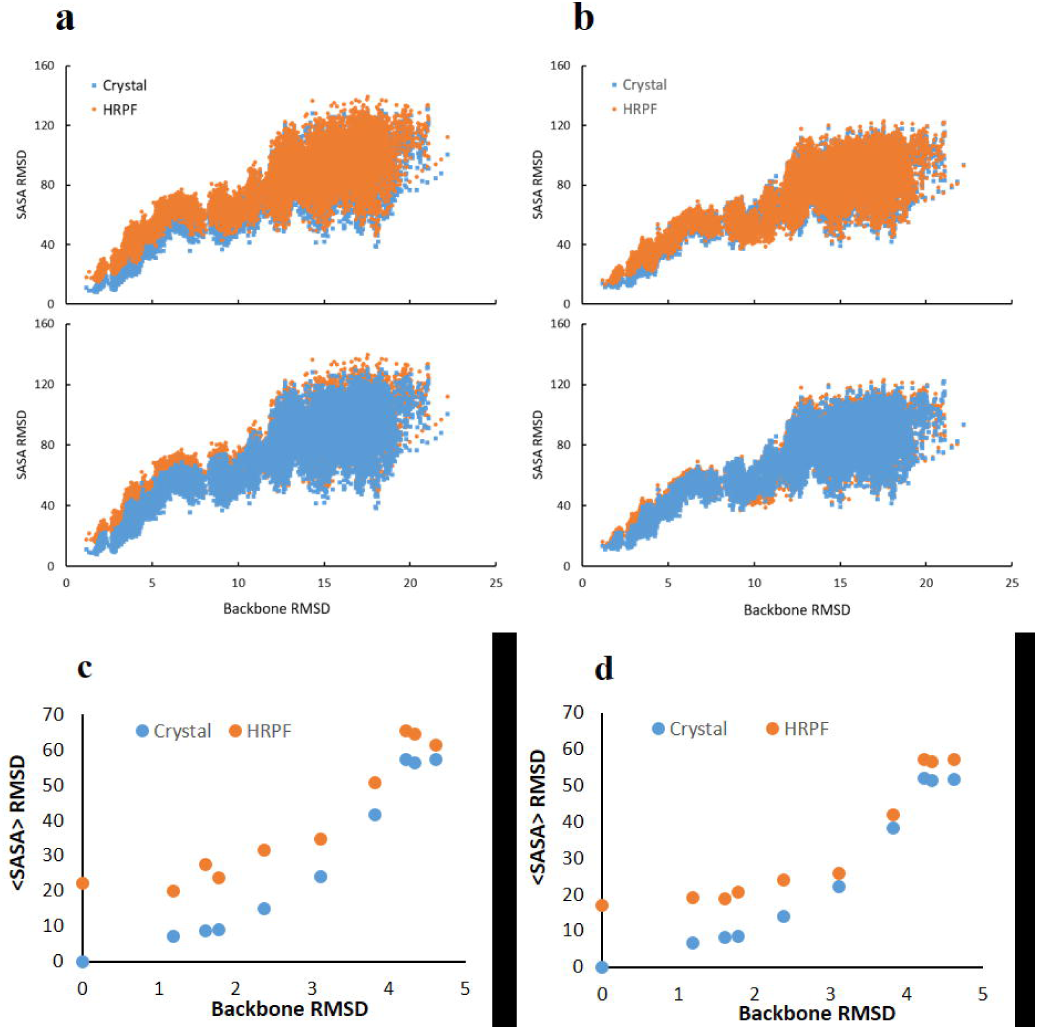
Evaluation of HR-HRPF derived <SASA> for assessing the quality of protein structural models. (**a**) SASA RMSD calculated on structure obtained from an unfolding MD simulation of lysozyme using crystallographic <SASA> and HR-HRPF derived <SASA> based on free amino acids normalization approach using only highly reactive amino acids as reference; (**b**) SASA RMSD calculated on structure obtained from an unfolding MD simulation of lysozyme using crystallographic <SASA> and HR-HRPF derived <SASA> based on the ratio of reactivity (native: denatured) approach as reference. Top and bottom panels of **a** and **b** show the same data, with different data series in front (**c**) SASA RMSD obtained from homology modeling analysis of lysozyme using crystallographic <SASA> and HR-HRPF derived <SASA> based on free amino acids normalization approach as reference. (**d**) SASA RMSD obtained from homology modeling analysis of lysozyme using crystallographic <SASA> and HR-HRPF derived <SASA> based on the ratio of reactivity (native: denatured) approach as reference.

While the HR-HRPF experimental <SASA> were able to differentiate MD models with good agreement to the native model in an MD unfolding experiment, many real-world applications of HR-HRPF data for computational modeling will rely upon the data to discriminate between homology models generated from various templates. To test the ability of HR-HRPF derived <SASA> to segregate good models from bad for such an approach, a set of homology models of lysozyme were generated using templates that yielded models with backbone RMSDs ranging from 0 Å to 5 Å (**Supplementary Table 3**). The SASA RMSD were calculated using the HR-HRPF derived <SASA> from each approach as reference and compared to the backbone RMSD (Fig. 5c,**d**). As these are static structures, the <SASA> RMSD is calculated from the SASA calculated from the X-ray crystal structure, rather than a structure relaxed using molecular dynamics simulations. This lack of solution phase-based relaxation leads to the apparent deviation between X-ray crystal-based RMSD SASA values (which measures static SASA in the crystalline lattice) and the solvent-based <SASA> measured by HRPF, resulting solely from how the “correct” <SASA> value is defined.

The approach based on the ratio of reactivity (native: denatured) provides more accurate <SASA> RMSD from the crystallographic data as compared to the approach based on normalization by free amino acids, which is consistent with the results obtained from the unfolding MD simulation study. Both approaches reveal a substantial increase in <SASA> deviation as backbone RMSD increases above 3 Å, which is very consistent with the discriminatory ability of both crystallographic and HR-HRPF <SASA> in MD simulations. The data indicate that both approaches can differentiate inaccurate or inappropriate computational models (with backbone RMSD higher than 4) from good computational models (with backbone RMSD smaller than 3) regardless of if the model is generated by MD or by homology modeling, demonstrating the potential of using the HR-HRPF derived <SASA> to experimentally evaluate molecular models of protein structure.

## DISCUSSION

By combining advances in radical dosimetry, accurate quantification of HRPF on individual amino acids by ETD fragmentation, and introducing multi-point FPOP HRPF, we are able to generate HRPF data with sufficient accuracy and robustness to quantitatively evaluate the relationships between HR-HRPF oxidation profiles and biophysical properties of the protein analytes. While previous results have strongly supported the effects of relative changes in HR-HRPF profiles, our results here show that sequence context effects on inherent amino acid reactivities, especially for amino acids with poor inherent reactivity, is a major confounding factor in the correlation of HR-HRPF profiles with absolute biophysical properties of the protein: namely, <SASA> values. By either minimizing these sequence context effects by solely relying upon highly reactive amino acids, or by normalizing for these sequence context effects by comparing HR-HRPF profiles of native and denatured structures, we can now measure <SASA> with reasonable accuracy.

Another potential confounding effect is that of tertiary and quaternary structure other than <SASA> impacting the apparent rate of oxidation (e.g. local scavenging of radicals by highly reactive amino acids, impacts of changes in amino acid “microenvironment” altering inherent reactivity). Based on the effects of sequence context identified here, it is expected that a similar pattern of effects would arise based on such higher-order structure effects: amino acids with poor inherent reactivity would show a greater influence from these effects, while amino acids with high inherent reactivity would be largely unaffected. However, as shown in Figure 3, once sequence effects are corrected for, there is no noticeable difference between amino acids with poor reactivity from those with high reactivity in either magnitude or direction of deviation from the <SASA>-based model, suggesting that the effects of higher-order structure other than <SASA> are minimal.

With the method described here, a sufficient number of <SASA> values may be obtained through multi-point HR-HRPF to allow for the accurate ranking of computational models. These results demonstrate the utility of HR-HRPF as a mass spectrometry-based method for generating molecular models that are thoroughly experimentally validated. The ability to generate experimentally validated molecular models using mass spectrometry, with its tolerance for sample impurity and heterogeneity and its low sample requirements, brings many important new structures within experimental reach. Our results indicate that the discriminatory power of this approach is strong, allowing for selection of only high quality models, and is sufficiently flexible to work for both MD trajectory models and homology models. The potential for such FPOP-based HR-HRPF approaches to quantitatively measure topography of integral membrane proteins should also be explored.

## METHODS

### Multi-point FPOP labeling

The two model proteins, lysozyme and myoglobin, used in this study were obtained from commercial sources (Sigma-Aldrich). 20 μM of each protein in 50 mM sodium phosphate buffer along with 17 mM glutamine as scavenger and 1 mM adenine as radical dosimeter were combined for FPOP experiment. FPOP labeling of each protein sample was performed at various final peroxide concentrations (10mM, 25mM, 50mM, and 100mM) by a 248 nm COMPexPro 102 high pulse energy excimer laser (Coherent) as described previously.^20^ The FPOP labeling in this experiment was set up using flow rate of 14.7 μL/min, with the excimer laser power adjusted to 100 mJ/pulse focused to a spot 10.5 mm^2^at a laser pulse repetition rate of 10 Hz, which provide calculated 20% unirradiated buffer region between irradiated segments to help prevent multiple irradiations per volume. The FPOP labeled products were collected in quench solution as previously reported to eliminate excess hydrogen peroxide and any long-lived secondary oxidants.^18, 20, 30^ Protein denaturation was performed right before FPOP irradiation by heating the protein sample to 90°C for 30 min for myoglobin sample as previously reported.^31^ For denaturation of lysozyme sample, TCEP was added to a final concentration of 500μM with heating at 90°C for 30 min before FPOP experiments. Adenine dosimeter UV absorbance at 260 nm was measured by a Thermo NanoDrop 2000c UV spectrophotometer (Thermo Fisher Scientific) to measure the effective radical dose delivered. Proteolytic digestion of oxidized protein was performed by addition of DTT to final concentration of 5mM followed by incubating at 65°C for 30 min. After cooling to room temperature, samples were digested overnight with sequencing grade trypsin (Promega) at 37°C at an enzyme: protein ratio of 1: 20 (wt/wt). The digested samples were stored at -20°C until LC-MS/MS analysis.

### LC-MS/MS analysis

All samples were analyzed on a Thermo Orbitrap Fusion (Thermo Fisher Scientific) coupled with the Ultimate 3000 Nano LC system (Dionex). Reverse-phase separation was accomplished using a 300 μm i.d. × 5 mm C18 PepMap 100 trap column with 5 μm particle size (Thermo Fisher Scientific) coupled with a 150 × 0.075 mm PepMap 100 C18 analytical column with 3 μm particle size (Thermo Fisher Scientific). Mobile-phase solvent A consisted of 0.1% formic acid in water, and mobile-phase B consisted of 0.1% formic acid in acetonitrile. The flow rate was set to 300 nL/min. A 45-min gradient was used to elute peptides (2% buffer B for 6 min, 2% to 40% buffer B within 24 min, 40% to 95% buffer B within 2 min, 95% buffer B for 2 min, 95% -2% buffer B within 1 min and 2% buffer B for 10 min).

The spray voltage was set to 2.6 kV. Full MS scan range was set to 300-2,000 *m/z* at a resolution of 60,000, and the automatic gain control (AGC) target was set to 2 × 10^5^. For the MS/MS scans, the resolution was set to 30,000, the precursor isolation width was 3 Da, ions were fragmented by both CID and ETD with the corresponding parent ion mass list. The CID normalized collision energy was 35%; the ETD-based precursor activation was carried out for 100 ms, with charge-state-dependent supplemental activation only applied on doubly-charged precursor ions.

### MS data analysis

Peptide sequences from both unoxidized controls and oxidized protein samples were screened using ByOnic (Protein Metrics, CA), searching against a database containing only the lysozyme or myoglobin sequence, with variable modifications set for common hydroxyl radical-mediated mass modifications.^2^All tandem mass spectra assignments and sites of oxidation were assigned manually due to peptide modification complexity. The tryptic digested peptides and their corresponding oxidation products were identified from the LC-MS runs. Identification and quantification of each oxidation site at the residue level was carried out using the ETD spectra collected from LC-MS/MS runs. Both peptide level and residue level quantitation were calculated as previously reported (**Supplementary Note 1**).^17, 32^

### <SASA> prediction model

The multi-point HR-HRPF regression was obtained by plotting the available radical concentration (measured by change in the absorbance of the UV radical dosimeter) versus the absolute level of oxidation for a given amino acid. The slope of the regression represents the radical response rate. Radical response rate of a residue *i* specifically measured for HR-HRPF normalized against the intrinsic reactivity of that free amino acid was calculated using the following equation modified from a similar approach reported previously:^23^

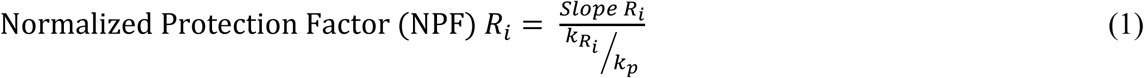

where Slope R_i_ is the slope of a specific residue *i* calculated from the multi-point HR-HRPF regression, and the k_Ri_/k_p_ is the residue-type specific intrinsic reactivity of residue *i* to hydroxyl radical on a relative scale to proline, to maintain with prior work in this area. The values of k_Ri_/k_p_ for 20 amino acids were compiled from previously reported radiolysis data in the literature.^24^

Sequence context study was performed by comparing the slope of the multi-point HR-HRPF regression from native folding protein to slope from fully denatured unfolding protein. The correlation study was conducted from the linear regression plotted using <SASA>/<SASA>_GXG_ versus NPF or <SASA>_N_/<SASA>_D_ versus Slope_N_/Slope_D_. The equation derived from the linear regression of the correlation study was then used to calculate <SASA> directly from HR-HRPF data.

### Molecular Modeling

Crystal structures of the globular proteins hen egg-white lysozyme (PDB ID: 2LYZ)^33^ and horse heart myoglobin (PDB ID: 1YMB)^34^ were used for the analysis. All waters of crystallization were removed from both the structures, along with any ions, such as the sulfate molecule present in 1YMB. The heme group in myoglobin was retained, as it is required for protein stability. All histidine residues were considered neutral and protonated only at the epsilon nitrogen. The N- and C-terminal residues were capped with acetyl (ACE) and *N*-methylamine (NME) groups, respectively. The proteins were minimized before solvating them in TIP3P water molecules, with a buffer size of 10 Å, resulting in the addition of 11299 and 9711 water molecules in the case of lysozyme and myoglobin, respectively. Counter ions (8 Cl^-^ ions) were added to neutralize the charge in lysozyme, using the tLEAP module of AMBER, no ions were required in the case of myoglobin.

Energy minimization was performed with the SANDER module of AMBER12^35^ with ff12SB protein force field parameters.^36^ Prior to solvation, the proteins were subjected to 3000 steps of steepest descent and 2000 steps conjugate gradient minimization, *in vacuo* to relieve any steric collisions. After solvation, the energy of the water and ions was minimized while keeping all protein atoms restrained (500 kcal/mol-Å^2^). The energy of the entire system was subjected to 5000 steps of steepest descent minimization, followed by 20000 steps of conjugate gradient minimization.

MD simulations of the energy-minimized systems were performed with the pmemd.cuda version of AMBER12.^37^All the bonds involving hydrogen atoms were constrained using the SHAKE algorithm,^38^ enabling an integration time step of 2 fs. Long-range electrostatic interactions were treated with the Particle-Mesh Ewald algorithm,^39^ with a long-range non-bonded interaction cut-off set to 10 Å. The systems were heated from 5 K to 300 K over a span of 50ps, under NVT conditions employing the Langevin thermostat.^40^ The simulations were then continued for 30 ns under NPT conditions with weak restraints on the Cα atoms of the protein (10 kcal/mol-Å^2^).

To generate an ensemble of partially-unfolded structures, the energy-minimized structure of unsolvated Lysozyme was heated during an MD simulation from 5 K to 1000 K over 10ns *in vacuo*. Snapshots were extracted from the simulation every ps. Over the course of the simulation, the all-atom RMSD of the conformations (with respect to the crystal structure) increased from 1.2 Å to 22.2 Å.

The per-residue solvent accessible surface area (SASA) was computed with the NACCESS program.^41^ Average SASA values (<SASA>) were computed from a total of 1000 snapshots extracted at 30 ps intervals from the solvated MD simulations.

The RMSD_SASA_ values were calculated using following equation,

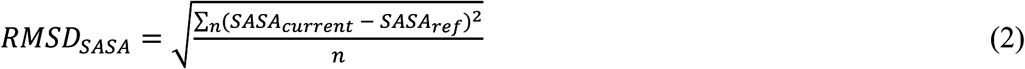

where SASA_current_ is the per-residue SASA value for each residue in the model (obtained from MD or from homology modeling) and SASA_ref_ is the SASA value for the same residue in the reference structure (either computed from the crystal structure or estimated from experimental HRPF data), and n is the total number of residues with SASA values.

Homology models for Lysozyme were generated using the SWISS-MODEL homology modeling server (swissmodel.expasy.org)^42^ for multiple template PDB structures with sequence identities that varied from 99% to as low as 37% with respect to Lysozyme (**Supplementary Table 3**). Templates were selected that had at least 90% sequence coverage to ensure plausible fold structures. The all-atom RMSD values (relative to Lysozyme, PDB ID: 2LYZ) ranged from as 1.2 to 4.6 Å.

## Acknowledgements

This research is supported by the National Institute of General Medical Sciences (1R01GM096049 and R01GM100058).

## Author Contributions Statement

B.X. performed all mass spectrometry experiments. B.X. and J.S.S. interpreted all mass spectrometry data and developed all experimental models for data normalization and conversion to SASA values. A.S. and R.J.W. performed all computational simulations and SASA calculations. All authors participated in the writing of this manuscript.

## Competing Financial Interests

J.S.S. discloses a significant ownership share of Photochem Technologies, LLC, a small company that is active in the area of hydroxyl radical protein footprinting instrument design.

